# Effects of variability in the amount and dispersion of within-plant herbivory on resistance- and tolerance-related responses in wild cotton

**DOI:** 10.1101/2025.07.14.664844

**Authors:** Víctor H. Ramírez-Delgado, Yeyson Briones-May, Juan Sánchez-Durán, Lucía Martín-Cacheda, Xoaquín Moreira, Ted C. J. Turlings, Luis Abdala-Roberts

**Author notes:** Universidad Nacional Autónoma de México Escuela Nacional de Estudios Superiores-Unidad Mérida Carretera Mérida-Tetiz Km. 4.5, Ucú, Yucatán 97357, Mexico (Luis Abdala-Roberts).

## Abstract

Herbivory triggers complex induced defensive responses in plants that may vary depending on the amount and dispersion of damage, the latter has received much less attention. We examined how wild cotton (*Gossypium hirsutum*) responds to different amounts of herbivory and how it is dispersed among the leaves (concentrated versus dispersed) using the specialist caterpillar *Alabama argillacea*. In a greenhouse experiment, plants underwent different herbivory treatments: no damage, low herbivory (two caterpillars, one on each of two leaves), high concentrated herbivory (four caterpillars, two on each of two leaves), and high dispersed herbivory (four caterpillars, each on one of four leaves). We measured extrafloral nectar (EFN), an indirect defense trait, and phenolic production, a direct chemical defense, as well as regrowth capacity (new leaf biomass two months after herbivory) as a tolerance proxy. To distinguish between local and systemic defense induction, we measured resistance-related responses on damaged and undamaged leaves. EFN was significantly and more consistently induced in damaged leaves (i.e., stronger local induction) in response to high dispersed damage. Phenolic compounds were not affected by any of the damage treatments. Additionally, wild cotton fully compensated for damage in terms of new leaf biomass under all herbivory treatments, except after high concentrated damage, suggesting higher costs incurred by this type of herbivory. These results highlight the importance of studying plant-induced defenses under varying herbivory conditions and suggest opposing effects on resistance-related (EFN, indirect) versus tolerance-related induced responses, which are known to affect tri-trophic interactions associated with wild cotton.

**Main Conclusion:** Variability in caterpillar damage to wild cotton leaves differentially shapes resistance- and tolerance-related responses; dispersed damage induces more extrafloral nectar production, and concentrated damage more strongly drives reductions in regrowth.

## Introduction

Herbivory has profound ecological and evolutionary impacts on plants (Strauss and Agrawal 1999; Karban 2011; Fritz and Simms 2012; Kant et al. 2015; Archibald et al. 2019; Adedayo et al. 2025). To mitigate the adverse effects, plants have evolved a wide array of defenses to either resist or tolerate herbivory (Leimu and Koricheva 2006, Núñez-Farfán et al. 2007; Fontes-Puebla and Bernal 2020). Direct resistance includes structural traits such as trichomes and chemical deterrents, including repellent or toxic secondary metabolites (Mauricio 2000; Núñez-Farfán et al. 2007; Endara et al 2023). Indirect resistance includes traits that attract natural enemies of herbivores by offering housing (e.g., domatia), signals that indicate the presence of herbivorous prey (volatile organic compounds), or food rewards such as extrafloral nectar (EFN) (Heil 2008; Kessler 2017). Tolerance, in contrast, is the plant’s ability to withstand or recover from herbivory, often through compensatory growth or resource reallocation, thereby minimizing fitness losses (Mauricio 2000; Núñez-Farfán et al. 2007; Salgado-Luarte et al. 2022; Liñán-Vigo and Núñez-Farfán 2024). Both resistance and tolerance strategies involve induced responses and resource reallocation, and they are often correlated (Strauss and Agrawal 1999; Leimu and Koricheva 2006; Oduor et al. 2011; Fornoni 2011) although this is not always the case (Leimu and Koricheva 2006; Carmona and Fornoni 2012; Kariñho-Betancour and Núñez-Farfán 2015).

The amount of herbivory is a key driver of plant resistance and tolerance strategies (War et al. 2012). Induced resistance mechanisms, such as the production of chemical defenses, are often herbivore dose-dependent, with higher damage levels frequently triggering stronger induced responses (Underwood 2000, 2010; Paniagua Montoya et al. 2024). In some cases, the responses are non-linear. For example, EFN sugar concentration, an indirect defense, may peak at intermediate damage levels and not increase upon further damage (Pimenta et al. 2025). Similar non-linear patterns have been reported for secondary metabolites (Karban and Baldwin 1997; Karban 2011; Eisenring et al. 2017). On the tolerance side, plants subjected to higher levels of damage are often unable to compensate fully, resulting in reduced growth or reproduction (Mundim et al. 2012; Tito et al. 2016; Quijano-Medina et al. 2019). These findings suggest that the costs of herbivory increase as damage levels exceed the plant’s capacity to reallocate resources for subsequent growth and reproduction (Núñez-Farfán et al. 2007). In some cases, overcompensation with increasing damage has been found for both vegetative and reproductive tissues (Paige 2018; Garcia and Eubanks 2019; Gols 2025). Understanding these damage-dependent responses is crucial for predicting plant fitness outcomes, the evolution of plant defensive strategies, and the coevolutionary dynamics between plants and herbivores.

The effects of varying amounts of herbivore damage on plants have been frequently studied, yet spatial patterning of damage among the leaves, i.e. how even (disperse) or uneven (concentrated) is distributed the damage, can be equally important in determining herbivory effects on plant responses and performance (The Herbivory Variability Network et al. 2023; Wetzel et al. 2023). The modular structure of plants and constraints on resource exchange among modules explain why they commonly better tolerate dispersed herbivory than concentrated herbivory (Marquis 1992, 1996; Mauricio et al. 1993). Indeed, when damage is concentrated, constraints to resource redistribution impair the recovery of affected modules, thereby reducing overall plant growth and reproduction (a phenomenon known as subcompensation). Less is known about the effects of variable degrees and patterns of herbivory on induced defense responses (but see: Ohnmeiss et al. 1997; Paniagua Montoya et al. 2024; Martín-Cacheda et al. 2025). Direct vascular connections that enable defense induction in undamaged leaves are linked to damaged ones via the exchange of metabolites as well as reallocation of resources among modules (Schittko and Baldwin 2003; Ferrieri et al. 2015; Eisenring et al. 2017). However, how different degrees of herbivory affect plant defense induction may differ for different plant traits and be dependent on the relative importance of systemic vs. local induction responses. On one hand, traits that are influenced by systemic responses are expected to be affected by physiological constraints that limit exchanges between different tissues (Schultz et al. 2013; Ferrieri et al. 2015). On the other hand, traits primarily induced locally may not exhibit such constraints, at least not in the short term, as their expression does not rely on inter-module resource reallocation; they occur within modules or parts of them (Marquis 1996). If this is the case, it is presumed that locally induced defenses are more strongly induced by concentrated damage in order to promote herbivore movement to other plant parts or neighboring plants (Marquis 1996). To date, very few studies have simultaneously tested how intensity and within-plant dispersion of herbivory affect resistance- and tolerance-related plant-induced responses. We here address this gap by studying resistance- and tolerance-related induced responses to herbivore damage in wild cotton (*Gossypium hirsutum* L.). Wild cotton occurs naturally in populations along the coast of northern Yucatan (Mexico), its likely center of origin and domestication (Wendel and Grover 2015). It is a myrmecophytic shrub that exhibits highly inducible direct (e.g., secondary metabolites) and indirect (e.g., EFN that attracts ants) resistance traits (Quijano-Medina et al. 2021; Briones-May et al. 2023; Mamin et al. 2025). It also appears to have tolerance mechanisms that mediate regrowth capacity in response to leaf damage, with a high amount of damage (e.g., 50% of total leaf area removal) leading to subcompensation (Quijano-Medina et al. 2019).

In our study, we inflicted different amounts and patterns of herbivore damage using the cotton specialist *Alabama argillacea*, which commonly occurs in the Yucatan cotton populations. In these populations, wild cotton plants exhibit a high degree of variability in the amount and distribution patterns of herbivore damage (Abdala-Roberts et al. 2019a), which has been shown to influence the induction of certain chemical defenses such as volatile organic compounds (Martín-Cacheda et al. 2025). In a greenhouse experiment, we exposed four-month-old plants to *A. argillacea* herbivory, varying the amount of damage as well as its within-plant dispersion. We then measured the levels of phenolic compounds and EFN, as well as leaf regrowth capacity. We hypothesized that plants would be better able to tolerate low than high damage (Quijano-Medina et al. 2019), more dispersed than concentrated damage, and, following from this, more strongly induce resistance-related traits in response to high vs. low damage and in response to concentrated vs. dispersed damage to mitigate the higher costs of high and concentrated damage. We further expected that resistance-related responses would be different for the two studied traits: EFN is largely induced locally (Abdala-Roberts et al. 2019b; Briones-May et al. 2023); therefore, it is less limited by physiological constraints and expected to be more strongly induced under concentrated damage. In contrast, phenolic compounds are induced systemically (Abdala-Roberts et al. 2019b), which may constrain their induction under concentrated damage.

## Methods

### Study species

Wild populations of *Gossypium hirsutum* L., commonly known as upland cotton, are distributed across southern Mexico, Central America, and the Caribbean, and are particularly common in the Yucatan Peninsula (Mexico) (Coppens d’Eeckenbrugge and Lacape 2014; Wendel and Grover 2015). In the Yucatan, populations are typically found in coastal shrublands, where leaf damage is primarily caused by grasshoppers and caterpillars (Abdala-Roberts et al. 2019a). Damage from these chewing insects is highly variable, showing up to a 3.5-fold difference among individuals within populations and up to a threefold difference in variability within plants (i.e., dispersion of damage among leaves on the same plant) (Abdala-Roberts et al. 2019a). Among the herbivores, the specialist caterpillar *A. argillacea* is particularly common and exhibits substantial variation in abundance across sites, ranging from rare to outbreak levels (L. Abdala-Roberts pers. obs.). Wild cotton defends itself against insect herbivores directly as well as indirectly (Hagenbucher et al. 2013). Direct defenses include toxic terpenoid aldehydes stored in pigmented glands (e.g., gossypol, heliocides; Opitz et al. 2008; Eisenring et al. 2017; Mamin et al. 2023), volatile organic compounds (also involved in plant-plant signaling; Grandi et al. 2024; Mamin et al. 2025), and phenolic compounds (Nix et al. 2017; Quijano-Medina et al. 2021). Phenolic compounds are particularly strongly induced in response to both natural and artificial herbivory in this species (e.g., Abdala-Roberts et al. 2019b; Quijano-Medina et al. 2021) and contribute to herbivore resistance (Nix et al. 2017; Quijano-Medina et al. 2024). In addition, wild cotton produces EFN in specialized nectaries, primarily located along the mid-vein on the ventral side of leaves, and EFN is also highly inducible in response to leaf damage (Abdala-Roberts et al. 2019b, Briones-May et al. 2023). However, we still know very little about the relative effects of herbivory amount and damage dispersion on the induction of secondary compounds such as phenolics, and on indirect defenses such as EFN (but see Martín-Cacheda et al. 2025).

### Plant material and propagation

In 2023, seeds were collected from seven adult *G. hirsutum* plants (hereafter referred to as “genotypes”) at a coastal scrubland site near Sisal, Yucatan, Mexico (21°11’45.3” N; 89°57’18.0” W). In February 2024, seeds were germinated, and seedlings were transplanted into 2-liter bags filled with a 1:2:1 mix of sand, native forest soil, and perlite. Plants were grown in a greenhouse at the Campus de Ciencias Biológicas y Agropecuarias (CCBA), Universidad Autónoma de Yucatán (20°52’00.6” N; 89°37’29.5” W), watered every other day with 300 ml of water, and used for experiments when they were four months old.

### Experimental Design

A total of 100 plants were placed in a greenhouse at the CCBA, spaced 90 cm apart. Plants were subjected to one of four herbivory damage treatments, with genotypes approximately evenly distributed across treatments. Damage was applied using clip cages containing a single third- or fourth-instar *A. argillacea* caterpillar. The treatments were as follows: (a) control: two clip cages with no larvae (one on leaf 2 and one on leaf 4, counting from the apical meristem) to control for clip cage effects; (b) low herbivory: two larvae, one on each of two leaves (leaves 2 and 4); (c) high concentrated herbivory: four larvae in total, with two larvae per leaf on leaves 2 and 4 (each larva placed in a separate clip cage, with cages on opposite sides of each leaf); and (d) high dispersed herbivory: four larvae total, one on each of four leaves (leaves 2, 3, 4, and 5). Larvae were placed on the plants for two consecutive nights (from 18:00 to 6:00 each day). We use nighttime to expose plants to larvae because the high daytime temperatures inside the greenhouse can cause the larvae to die. Under this setup, plants of the low-damage treatment experienced approximately half the damage compared to the high-damage treatments. Whereas both high-damage treatments resulted in similar total leaf area consumed, allowing us to test for the effects of damage dispersion while holding overall damage constant. To verify that these treatments achieved the intended herbivory levels, we photographed leaves 2–5 of all plants after the treatment period. Using ImageJ (version 1.59, Schneider et al. 2012), we measured total and damaged leaf area and calculated the percentage of herbivory from these four leaves. As expected, the two high-damage treatments did not differ significantly in total damage in leaves 2-5, but both resulted in approximately twice the damage observed in the low-damage treatment (Fig. S1, supplementary material).

### EFN volume and sugar quantification

EFN is rapidly induced after herbivory (within 12–48 h; Wäckers and Bonifay 2004; Abdala-Roberts et al. 2019b; Briones-May et al. 2023). Accordingly, we collected EFN from 98 plants the morning following the second night of larval exposure, between 6:00 and 8:00, using 5-μL capillary tubes (micropipettes Blaubrand® intraMARK, white color code, Germany). For each plant, separate samples were taken from leaves 1 and 2 (damaged and undamaged, respectively) to distinguish between local and systemic induced responses (Abdala-Roberts et al. 2019b; Briones-May et al. 2023). All nectaries were reset 24 h before sampling by removing existing nectar with cotton wool. After measuring EFN volume with capillary tubes, we assessed sugar concentration using a refractometer (Atago Master, Germany), recorded in ° Brix. Total sugar production per leaf was then estimated by multiplying nectar volume by sugar concentration, assuming 1 ° Brix equals 1 g of sugar per 100 g of solution (Kimball 1991).

### Quantification of phenolic compounds

All plants remained in the greenhouse after EFN sampling, and a subsample of 50 plants was used for phenolic analysis (randomly selected within each treatment and evenly distributed across treatments and genotypes). These plants were sampled one week after the herbivory treatments by harvesting leaves 1 and 2 (the youngest leaves). Collected leaves were immediately dried at 40 °C for three days (Moreira et al. 2025). This sampling procedure was chosen because the induction of phenolic compounds induced responses is typically slower, with significant effects on preexisting leaves reported to emerge between 72 h and one week after damage (Abdala-Roberts et al. 2019b; Quijano-Medina et al. 2023). Total phenolic content was determined using the Folin–Ciocalteu colorimetric method (Adegbusi et al. 2022). A standard curve was constructed using tannic acid, and results are reported as milligrams of tannic acid equivalents (TAE) per g of dry weight (DW).

### Regrowth responses

The remaining 50 plants that were not used for phenolic quantification were assessed for their regrowth capacity as a proxy for tolerance to herbivory. These plants were kept in the greenhouse for an additional two months, after which all newly produced leaves (post-treatment) were harvested and dried at 45 °C for one week. The total DW of the new leaves was then measured using an analytical balance (±0.1 mg, FA2004, Liberty, USA) to evaluate treatment effects on regrowth in terms of leaf biomass (Quijano-Medina et al. 2019).

### Statistical analyses

We tested the effect of damage treatment (fixed, four levels) on nectar volume and total sugar production using generalized linear mixed models (GLMMs) with a Tweedie distribution and a log link function. The Tweedie distribution provided a good fit for the EFN data, which exhibited a right-skewed distribution with zeros (Hasan and Dunn 2015). In addition, we employed linear mixed models (LMMs) for phenolic compound concentrations and the DW of new leaves. All models included plant height as a covariate to account for variability in plant size that could influence induced responses and regrowth capacity, and plant genotype (source mother plant) as a random factor to account for background variation due to genetic or maternal effects.

We performed all statistical analyses using R 4.4.2. (R Core Team 2024) with the following packages: “glmmTMB” to fit GLMMs (Brooks et al. 2017), “lme4” to fit LMMs (Bates et al. 2015), “DHARMa” to evaluate model fit (Hartig 2024), and “car” to test the main effects in GLMMs using Wald χ^2^ tests (Fox and Weisberg 2019). In addition, we used “lmerTest” to compute F- and P-values for the main effects in LMMs (Kuznetsova et al. 2017), “emmeans” to perform Tukey pairwise contrasts between treatment groups and to extract back-transformed estimated marginal means (EMMs) with 95% confidence intervals (CIs) from the models (Lenth 2024), and “ggplot2” to visualize the results (Wickham 2016). Throughout the text, results are reported as EMM [2.5% lower CI, 97.5% upper CI].

## Results

### Resistance-related induced defenses

#### Extrafloral nectar

We found no significant effect of the damage treatment on EFN volume or sugar content produced by leaf 1 (undamaged, i.e., systemic induction) (Table 1; Fig. 1a, b). However, there was a significant effect of the damage treatment on both nectar volume and concentration for leaf 2 (damaged, i.e., local induction) (Table 1). This effect was most evident for the volume produced after high dispersed damage treatment (0.598 µL [0.408, 0.876]), followed by the low damage treatment (0.423 µL [0.281, 0.636]), which did not differ significantly from the former (Fig. 1c). These two treatments differed significantly from both the control (0.17 µL [0.105, 0.276]) and the high concentrated treatment (0.144 µL [0.084, 0.244]), with the latter not differing from the control (Fig. 1c). Regarding leaf 2 sugar content, high dispersed damage produced the highest mean value (0.375 µg [0.292, 0.482]) and differed significantly from all other groups, including the control (0.158 µg [0.11, 0.228]) (Fig. 1d). The other damage treatments (low damage: 0.214 µg [0.154, 0.297]; high concentrated damage: 0.139 µg [0.091, 0.211]) did not differ from each other or the control (Fig. 1d).

**Fig. 1.**
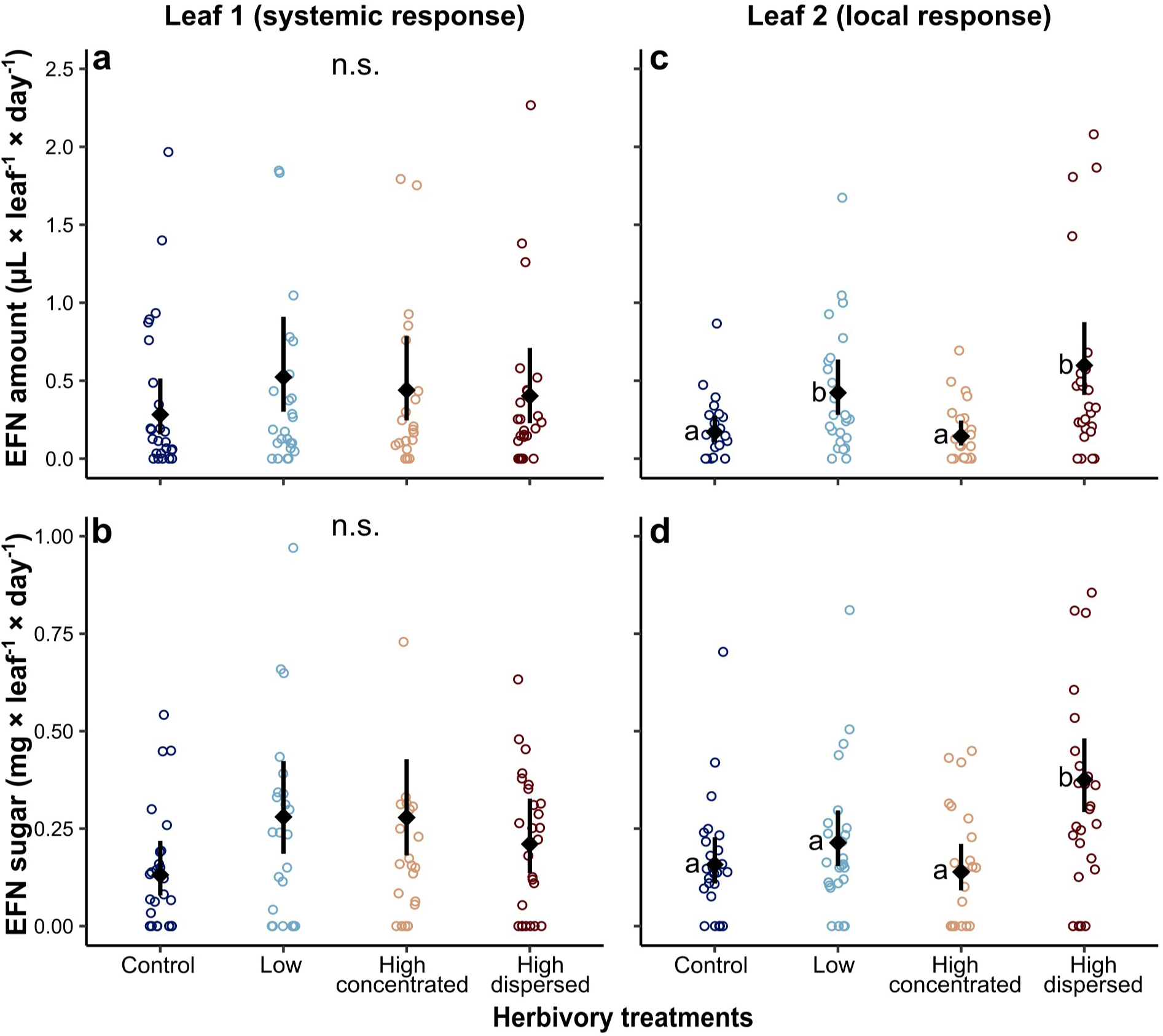
Effects of damage amount and dispersion on extrafloral nectar (EFN) production, in terms of nectar volume and total sugar content, in wild cotton (*Gossypium hirsutum*). Panels show: (a) leaf 1 (undamaged, reflecting systemic induction) and (b) leaf 2 (damaged, reflecting local induction) for EFN volume (µl) produced and total sugars (mg) in response to damage treatments. Colored circles represent individual plant values for each herbivory treatment (dark blue: control, blue: low, yellow: high concentrated, dark red: high dispersed herbivore). Black squares and error bars indicate estimated marginal means and 95% confidence intervals from general linear mixed models. Different letters indicate significant differences among treatments (Tukey post-hoc tests, *P* < 0.05); “n.s.” denotes non-significant differences.

**Table 1.** Statistical analysis of plant defense responses in relation to damage treatments and plant height. Results from general linear models for induced extrafloral nectar (EFN) volume and sugar production, and from linear mixed models for induced phenolic compounds and tolerance (dry weight of new leaves). Significant *p*-values (*p* < 0.05) are shown in bold. The table presents test statistics, degrees of freedom, and *p*-values for each effect (damage treatment or plant height), measured at two leaf levels when applicable.

| Effect | Response variable |  |  |  |  |  |  |
| --- | --- | --- | --- | --- | --- | --- | --- |
|  | EFN volume |  | Total sugar |  | Total phenolics |  | Tolerance |
|  | Leaf 1 | Leaf 2 | Leaf 1 | Leaf 2 | Leaf 1 | Leaf 2 |  |
| Damage treatment | $\chi^2 = 3.54$;<br>df = 3;<br>$p = 0.3158$ | $\chi^2 = 32.69$;<br>df = 3;<br>$p < \mathbf{0.0001}$ | $\chi^2 = 7.76$;<br>df = 3;<br>$p = 0.0512$ | $\chi^2 = 21.84$;<br>df = 3;<br>$p < \mathbf{0.0001}$ | F = 1.87;<br>df = 3, 40.8;<br>$p = 0.1491$ | F = 0.56;<br>df = 3, 40.08;<br>$p = 0.6427$ | F = 4.53;<br>df = 3, 40.07;<br>$p = \mathbf{0.0079}$ |
| Plant height | $\chi^2 = 5.26$;<br>df = 1;<br>$p = \mathbf{0.0218}$ | $\chi^2 = 5.77$;<br>df = 1;<br>$p = \mathbf{0.0163}$ | $\chi^2 = 0.70$;<br>df = 1;<br>$p = 0.4037$ | $\chi^2 = 4.19$;<br>df = 1;<br>$p = \mathbf{0.0227}$ | F = 0.10;<br>df = 1,<br>43.07;<br>$p = 0.7471$ | F = 0.24;<br>df = 1, 44.78;<br>$p = 0.6257$ | F = 2.68;<br>df = 1, 42.84;<br>$p = 0.1085$ |

#### Phenolic content

Although there was a trend toward higher induction under the high dispersed damage treatment (Fig. 2), we found no significant effect of the damage treatment on phenolic content for either leaf 1 or leaf 2 (Table 1).

**Fig. 2.**
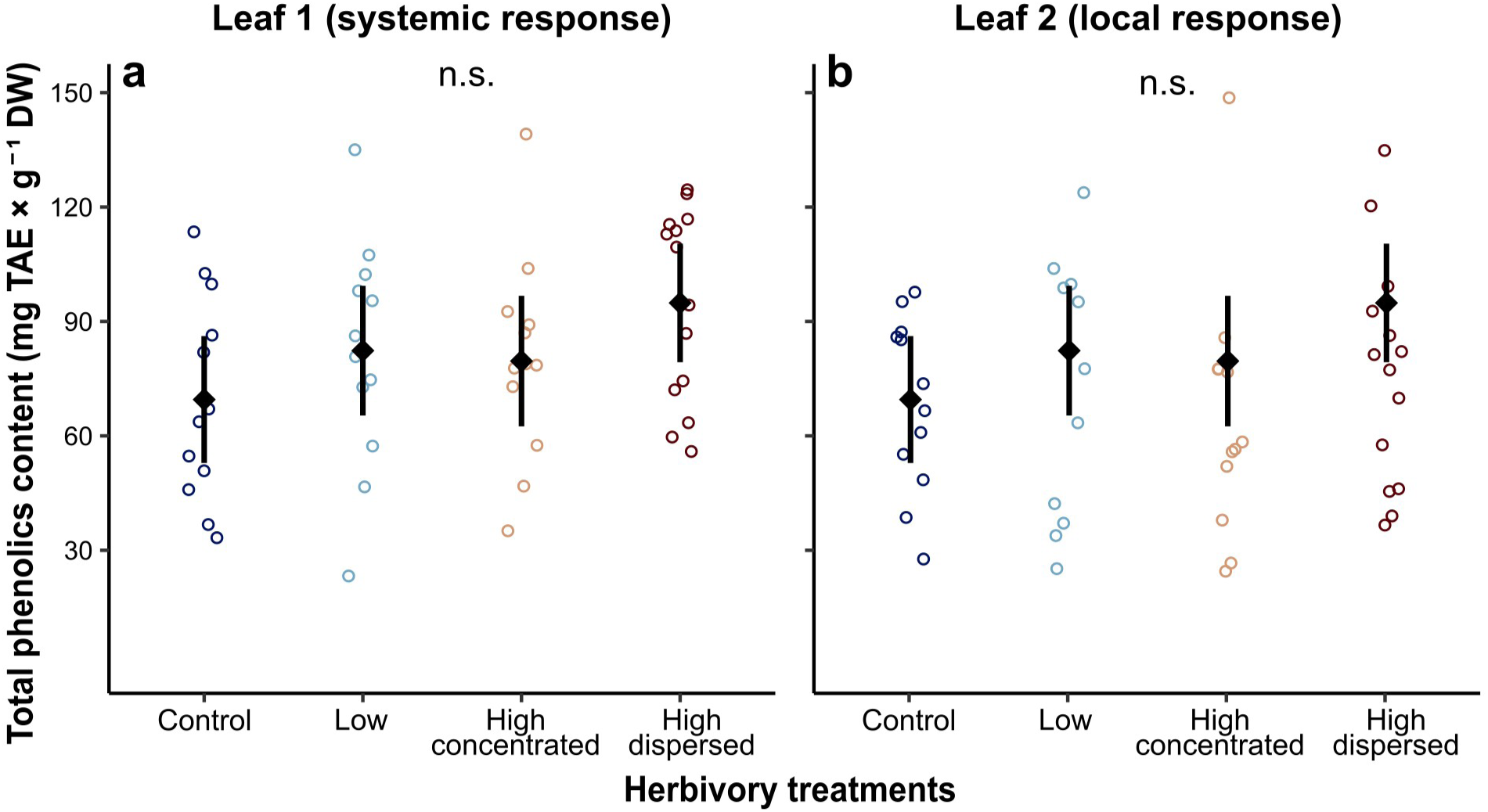
Effects of damage amount and dispersion on phenolics compound production in wild cotton (*Gossypium hirsutum*). Panels show: (a) leaf 1 (systemic response) and (b) leaf 2 (local response), with phenolic concentrations expressed as tannic acid equivalents (TAE) per gram of leaf dry weight (DW). Colored circles represent individual plant values for each herbivory treatment (dark blue: control, blue: low, yellow: high concentrated, dark red: high dispersed herbivore). Black squares and error bars indicate estimated marginal means and 95% confidence intervals from linear mixed models. There were no significant differences among treatments, as indicated by “n.s.”

### Tolerance-related defensive responses

We found a significant effect of the damage treatment on the DW of new leaves produced after damage (Table 1). Group comparisons showed that plants subjected to high concentrated damage (1.38 g [1.16, 1.6]) were the only treatment group with a significantly lower mean DW than control plants (1.87 g [1.67, 2.07]) (Table 1, Fig. 2). Despite a trend, high concentrated damage did not differ significantly from the other damage treatments (low damage: 1.6 g [1.4, 1.8]; high dispersed damage: 1.6 g [1.4, 1.8]), and these two treatments also did not differ from each other (Fig. 3).

**Fig. 3.**
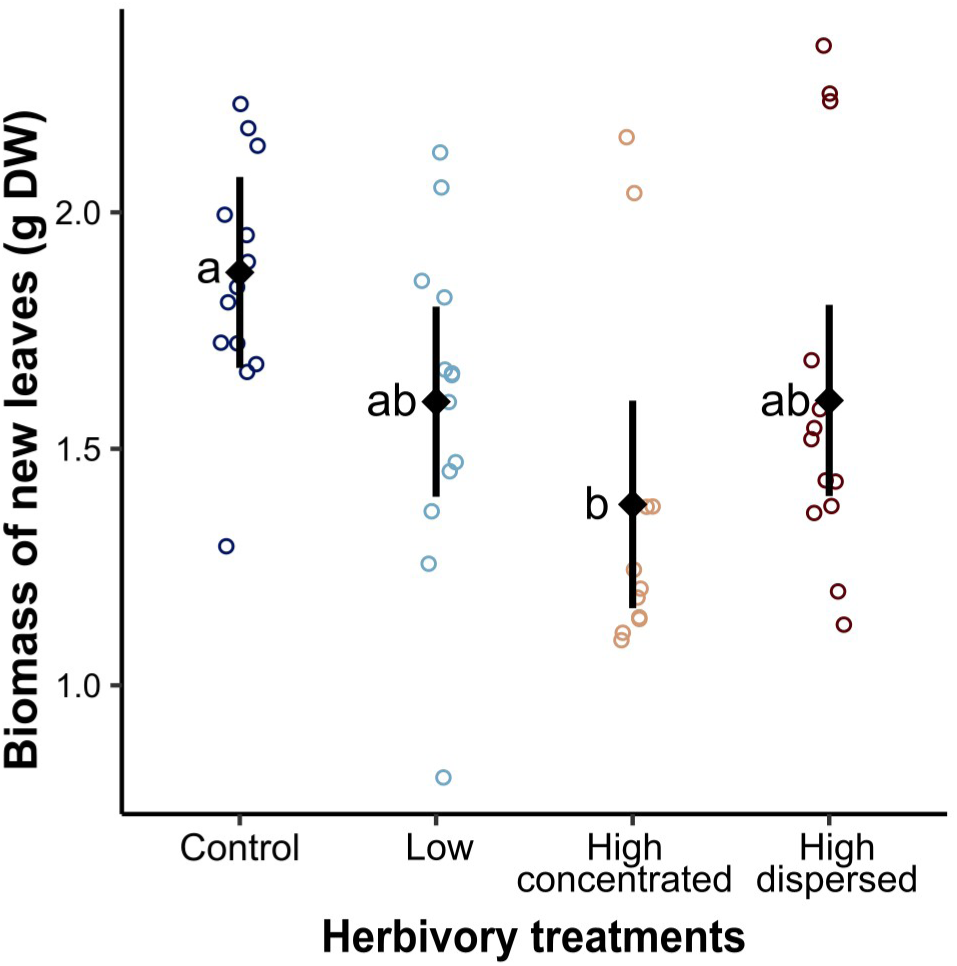
Effects of herbivory amount and within-plant variability on the biomass of newly developed leaves, measured as grams of dry weight (DW) after the herbivory treatments were applied. Colored circles represent individual plant values for each treatment (dark blue: control, blue: low, yellow: high concentrated, dark red: high dispersed herbivore). Black squares and error bars indicate estimated marginal mean and 95% confidence intervals from linear mixed models. Different letters indicate significant differences among treatments (Tukey post-hoc test, *P* < 0.01).

## Discussion

Our study shows that the characteristics of herbivory, namely its amount and its spatial dispersion among the leaves, differentially affect resistance- and tolerance-related defensive responses in wild cotton. Consistent with our predictions, we found that regrowth capacity was mainly compromised under concentrated damage, suggesting higher costs associated with this herbivory pattern and highlighting the role of within-plant herbivore variability as a key driver of tolerance responses in cotton. Contrary to our expectations, EFN induction, particularly sugar content, was more strongly triggered by dispersed damage (with a similar trend for EFN volume, albeit comparable to that observed under low damage), while phenolic content was not significantly increased in response to any damage treatment. Therefore, although concentrated damage may have a greater negative impact on fitness, it did not lead to stronger induction of resistance-related traits that might mitigate these costs. Overall, these findings emphasize the importance of considering variation in herbivory and accounting for spatial patterns of damage when investigating plant defensive strategies and their associated induced responses. The significant reduction in new leaf DW observed in plants subjected to concentrated damage supports the hypothesis that concentrated herbivory imposes greater costs, possibly through physiological constraints, than dispersed damage. This finding aligns with previous research on the effects of plant modularity on plant tolerance responses, which emphasizes constraints on resource redistribution and regrowth capacity (Marquis 1992; Mauricio et al. 1993; Wetzel et al. 2023). Although the four-month-old cotton plants used in our experiment had not yet started branching, our results suggest that these constraints operate even among leaves within the same stem. Because our concentrated herbivory treatment targeted alternate leaves, these constraints could also act at the level of orthostichies, with the affected orthostichy contributing less to regrowth due to reduced photosynthetic capacity (Price and Hutchings 1992; Marquis 1996). Such limitations likely involve direct disruption of vascular integrity, e.g., impairing water and nutrient transport by creating non-functional xylem conduits and compromising phloem pathways (Brodersen and McElrone 2013). Additionally, herbivory reduces the photosynthetic capacity of wounded modules (Schultz et al. 2013). Together, these processes can significantly reduce the viability of heavily damaged tissues and impair the regrowth capacity of individual plants. In our study, herbivory levels were relatively low (20-30% leaf area removed for 2 to 4, out of 10-12 leaves per plant), compared to a previous study that found regrowth deficits in plants experiencing moderate to high mechanical damage (e.g., 50% of total leaf area removed in younger plants with 8–10 leaves; Quijano-Medina et al. 2019). This result suggests that concentrated damage can pose a significant challenge to regrowth in wild cotton, even at low to moderate levels. Mechanisms such as resource reallocation (e.g., exchange of soluble sugars between leaves and roots; Quijano-Medina et al. 2019) and increased photosynthetic activity may not be sufficient to offset the negative impacts of herbivory under these conditions.

Consistent with previous work, EFN was more strongly induced locally, that is, when analyzing damaged leaves, and showed no or only weak systemic effects (Abdala-Roberts et al. 2019b; Briones-May et al. 2023). When comparing treatment groups, particularly regarding nectar sugar amounts, the high dispersed herbivory treatment induced the strongest EFN response and differed significantly from the control. While this finding contradicts our initial prediction (i.e., stronger resistance-related defense induction under concentrated damage), it still highlights the importance of herbivory variability—rather than total herbivory amount, at least within the levels tested—in shaping EFN responses. We hypothesize that signaling pathways associated with EFN induction may be more effectively activated when damage is spread across multiple modules, as risk signals may plateau within individual leaves but accumulate across damaged modules (de Bruxelles and Roberts 2001). At the same time, plants may limit EFN induction in heavily damaged modules to allocate resources toward other resistance-related traits, reflecting well-documented tradeoffs between defense strategies in plants (Moles et al. 2013; Eichenberg et al. 2015). In turn, the observed EFN responses to within-plant variability in herbivore damage may have important downstream consequences for ant-plant interactions (Llandres et al. 2019; Vázquez-Barrios et al. 2021; Reyes-Hernández et al. 2022), which could explain EFN induction patterns. Supporting this, ongoing work shows that dispersed damage leads to a greater number of secreting nectaries (a response not measured here, presumably due to local induction), which increases ant recruitment and enhances herbivore attack rates (Ramírez-Delgado et al. under review). Thus, both increased EFN secretion per nectary (as shown here) and whole-plant level induction (perhaps more importantly) confirm greater EFN investment in response to dispersed damage, mediating indirect defense via ant recruitment. These findings underscore that future studies should consider the effects of within-plant herbivory variability and its underlying mechanisms in upland cotton, as well as in other *Gossypium* species (e.g., see Wäckers et al. 2001; Rudgers et al. 2004) and other EFN-bearing myrmecophytic species.

The lack of a significant treatment effect on phenolic compounds levels was unexpected. Previous studies with wild cotton using artificial damage (e.g., mechanical wounding with *Spodoptera frugiperda* oral secretions or *S. frugiperda*-only damage) reported significant induction of phenolic compounds in preexisting leaves within 72 h and up to one week after damage onset (Abdala-Roberts et al. 2019b; Quijano-Medina et al. 2023). In addition, another study found significant induction of phenolic compounds one week after damage; however, these responses were generally weaker than those observed in newly developed leaves collected five weeks post-damage (Quijano-Medina et al. 2021), suggesting that systemic phenolic responses in new tissues play a more important role in induced resistance. Earlier studies with cultivated cotton and wild cotton (Usha Rani and Pratyusha 2013; Dixit et al. 2017), using generalist lepidopteran herbivores (*Spodoptera litura* and *Helicoverpa armigera*), also reported significant induction of phenolic compounds one to four days after damage onset, along with deleterious effects on herbivores. These findings suggest that phenolic compounds are generally highly inducible in cotton across a range of experimental approaches, herbivore species, and damage types (natural vs. artificial), and across both domesticated and wild cotton. However, we cannot rule out the possibility that peak induction occurred earlier than our sampling time (e.g., between 48 and 72 h post-damage) or that systemic induction was stronger in newly developed leaves —an effect also observed for terpenoid aldehydes (Opitz et al. 2008; Eisering et al. 2017, 2018.). Another possibility is that *A. argillacea*, as a specialist, suppresses phenolic compound production. Herbivore-mediated suppression of plant defenses is a known phenomenon (reviewed by Ali and Agrawal 2012). It has been documented in both specialist systems, e.g., the suppression of glucosinolates in *Brassica oleracea* (Karssemeijer et al. 2022), and generalist systems, such as the suppression of green leaf volatiles in *Zea mays* (Jones et al. 2019), and jasmonic acid-mediated defenses in tomato by *Aculops lycopersici* (Glas et al. 2014), and *Arabidopsis thaliana* (Chen et al. 2019). Consistent with our findings, ongoing work also shows non-significant induction by this insect across two ontogenetic stages of wild cotton, i.e., one month- vs four-month-old plants (Briones-May et al. under review). Further research is needed to test if *A. argillacea* is indeed able to suppress defenses in cotton, for instance by assessing defensive gene expression levels in response to varying (and potentially higher) caterpillar loads and comparing the responses with those induced by other caterpillar species.

### Gaps in research and directions

Although our study provides novel insights into how herbivory patterns influence defense allocation, several questions remain. Future research should explore the temporal dynamics of these responses, as defense induction and tolerance mechanisms may operate on different timescales. Additionally, investigating whether these patterns hold across different herbivore species, particularly when comparing specialists to generalists, would further illuminate the context dependency of defense allocation strategies under varying herbivory scenarios. Examining the molecular mechanisms underlying differential responses to concentrated versus dispersed damage could also provide deeper insights into the signaling pathways that regulate resistance- and tolerance-related responses. Lastly, an important gap to address is the measurement of other types of chemical and structural defenses in cotton, such as terpenoid aldehydes and trichome density (pubescence).

## Conclusion

Our study demonstrates that *G. hirsutum* tailors its defensive strategies based on the spatial dispersion of herbivory within individual plants, rather than simply responding to total damage levels. These results suggest that damage characteristics both enable adaptive responses under different threat scenarios and constrain defensive options. These findings highlight the importance of examining herbivory variability alongside mean damage levels when studying induced defenses. Understanding how plants have evolved to detect and respond to complex herbivory patterns carries broader ecological and evolutionary implications for plant-herbivore interactions.

## Data archiving

The data and analysis scripts for this study have been deposited in the Zenodo repository and are available at [DOI]

