## Supplemental Fig. S1 for "Effects of variability in the amount and dispersion of within-plant herbivory on resistance- and tolerance-related responses in wild cotton"

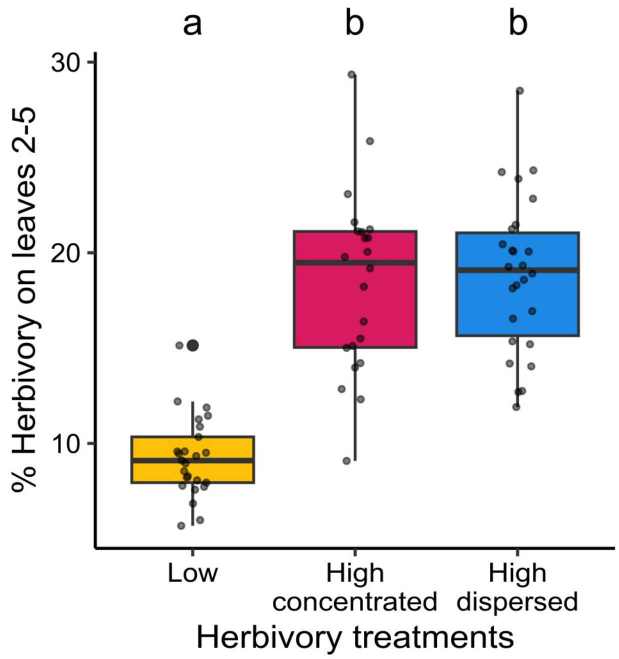


**Fig. S1** Boxplots show the percentage of tissue removed on leaves 2–5 for each herbivory treatment group. Box elements represent median (center line), interquartile range (box), and full data range (whiskers). Individual data points are overlaid to show the distribution of values. One-way analysis of variance revealed significant differences among treatments ($F_{2,70} = 51.24$, $P < 0.0001$). Post-hoc Tukey's HSD tests showed that herbivory damage differed significantly between the low and both high concentrated ($P < 0.0001$) and high dispersed damage ($P < 0.0001$) treatments, but there was no significant difference between the high concentrated and high dispersed herbivore damage treatments ($P = 0.9485$). Different letters indicate these pairwise differences.
